# Interspecies activation correlations reveal functional correspondences between marmoset and human brain areas

**DOI:** 10.1101/2021.02.09.430509

**Authors:** Yuki Hori, Justine C. Cléry, David J. Schaeffer, Ravi S. Menon, Stefan Everling

## Abstract

The common marmoset has enormous promise as a nonhuman primate model of human brain functions. While resting-state functional magnetic resonance imaging (fMRI) has provided evidence for a similar organization of marmoset and human cortices, the technique cannot be used to map the functional correspondences of brain regions between species. This limitation can be overcome by movie-driven fMRI (md-fMRI), which has become a popular tool for non-invasively mapping the neural patterns generated by rich and naturalistic stimulation. Here, we used md-fMRI in marmosets and humans to identify whole-brain functional correspondences between the two primate species. In particular, we describe functional correlates for the well-known human face, body, and scene patches in marmosets. We find that these networks have a similar organization in both species, suggesting a largely conserved organization of higher-order visual areas between New World marmoset monkeys and humans. However, while face patches in humans and marmosets were activated by marmoset faces, only human face patches responded to the faces of other animals. Together, the results demonstrate that md-fMRI is a powerful tool for interspecies functional mapping and characterization of higher-order visual functions.

## Introduction

The common marmoset has become an important nonhuman primate model for bridging the translational gap between rodents and humans. Marmosets have a lissencephalic cortex, like rodents, but as primates they possess the structural and functional brain architecture that supports elaborated behaviors such as pro-social interaction (Ferrari and Digby, 1996; Huang et al., 2020; Miller et al., 2016; Saito, 2015; Yokoyama and Onoe, 2015), complex vocal communication (Choi et al., 2015; Miller et al., 2015; Sadagopan et al., 2015; Takahashi et al., 2016), and saccadic eye movements (Chen et al., 2020; Ma et al., 2020; Mitchell et al., 2014; Selvanayagam et al., 2019). This, paired with a high reproductive power, small size, and fast maturation rate, make this non-human primate (NHP) species particularly interesting for neuroscience.

Functional magnetic resonance imaging (fMRI) plays an important role in identifying the functional architecture of marmoset cortex (Belcher et al., 2013; Hori et al., 2020a; Schaeffer et al., 2019c) and in establishing brain homologies between primate species (Ghahremani et al., 2016; Hori et al., 2020b; Schaeffer et al., 2019c). In particular resting-state fMRI has been used (i) to identify homologous large-scale brain networks between marmosets and humans (Ghahremani et al., 2016; Hori et al., 2020a; Schaeffer et al., 2019c), (ii) to define functional boundaries based on intrinsic functional connectivity (Schaeffer et al., 2019b, 2019a), and (iii) to use functional connectivity “fingerprints” of brain areas to establish homologies between marmosets, rodents, and humans (Schaeffer et al., 2020a). Because resting state patterns are state-agnostic and spontaneous, however, this technique cannot be used to map the functional correspondences of interspecies BOLD fluctuations over time. Task-based fMRI is better suited for mapping stimuli-driven fluctuations across species and indeed a few studies have used task-based fMRI in awake marmosets to identify areas related to specific functions (e.g. visuo-saccadic orienting (Schaeffer et al., 2019d), processing of faces and bodies (Hung et al., 2015; Schaeffer et al., 2020b), looming and receding visual stimuli (Cléry et al., 2020, Neuroimage), and tactile processing (Clery et al., 2020). A major drawback of task-based fMRI is that compliance is often poor in NHPs and each task can only reveal the limited set of functional activations for which it was designed.

These limitations can be overcome by employing movie stimuli which provide rich and naturalistic stimulation. Human studies have shown that movie-driven fMRI (md-fMRI) responses are highly selective between brain regions, engage many brain regions, and are highly reliable between subjects (Hasson et al., 2010, 2008; Lerner et al., 2011; Naci et al., 2014). Functional correspondences between species can be directly tested by the interspecies activity correlation (ISAC) method which uses the md-fMRI time course in a seed region in one species to identify functional correspondences across the cortex of the other species (Mantini et al., 2012). This technique has been successfully employed to identify functional correspondences (analogies) between human and macaque cortical areas, but this powerful mapping technique has yet to be applied to the marmoset brain.

Here we used md-fMRI in marmosets and humans to establish functional correspondences between cortical areas across the brain in the two species. We focused our analysis on identifying analogies of the well-known human face-, body-, and scene-selective networks in the marmoset brain. Not only do these networks play pivotal roles in primate visual perception, face-selective areas have already been described by a few task-based fMRI studies in marmosets, therefore providing an independent validation of the results.

## Results

Thirteen human participants and four marmoset monkeys freely watched a 15 min naturalistic movie with no sound. The movie was selected to keep marmosets interested and showed a fictitious day of a marmoset living in and navigating through an urban environment. Aside from marmosets (present in ~57% of the movie), a variety of other species (humans, owls, capybaras, dogs, cats, pigeons, roosters, frogs, ants) are also present in some parts (~27%). The movie was presented once to each human participant and 3-4 times to each monkey on separate days.

### Inter-scan variability of the stimulus driven activity

We first identified the cortical brain regions with consistent md-fMRI activity among scans to remove the low reproducible areas from the subsequent analysis. To this end, we calculated inter-scan correlation maps for the human and marmoset participants. We found high correlations between scans in occipital, temporal, and parietal areas in both species (Fig. 1A-F; please see the Suppl. Fig. 1A-D for the right hemisphere). Unlike humans, marmosets also showed high correlations in frontal and prefrontal areas. These findings are very similar to a previous human and macaque monkey study (Mantini et al., 2012). A difference from this previous study is a lack of correlations in human auditory areas because our subjects watched the movie with no sound.

**Fig. 1.**
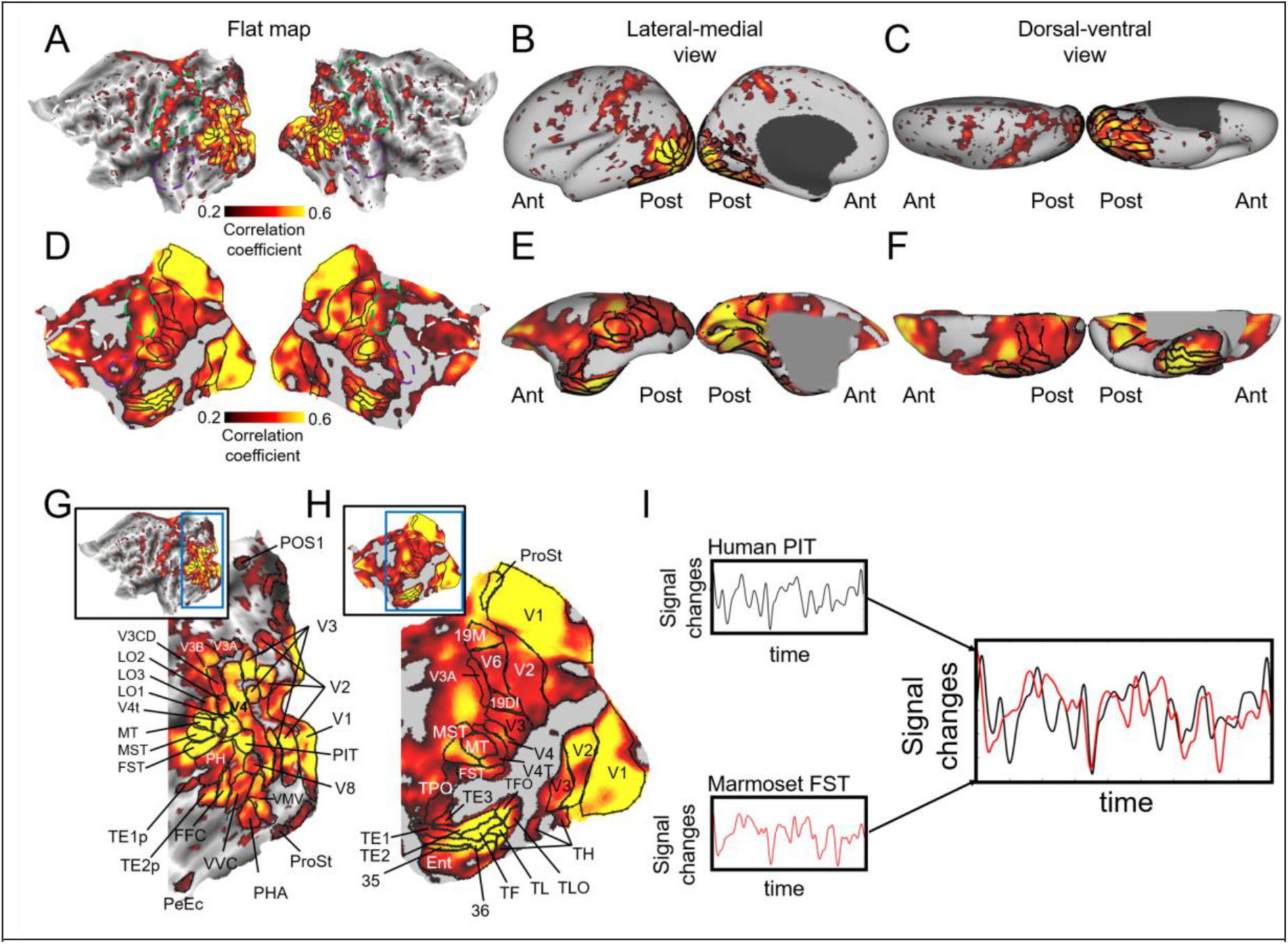
Inter-scan correlation map of brain activity during movie viewing. Spatial maps of correlated brain activity across 13 scans (i.e. 13 human subjects) and across 14 scans (4 marmoset subjects), mapped on each flattened cortex (A, D) and left cortical surface (lateral-medial view (B, E) and dorsal-ventral view (C, F)). Approximate locations of parietal, auditory and frontal regions are indicated green, purple, and white dashed lines on the flat maps, respectively. The volume-of-interests (VOIs) in visual-related areas were manually created based on the multi-modal cortical parcellation atlas (Glasser et al., 2016) for humans and Paxinos atlas for marmosets (Liu et al., 2021), so as not to include the low correlation areas among scans (r < 0.2) (G for humans, and H for marmosets). Then, the time courses were extracted from two VOIs (e.g. PIT in humans and FST in marmosets) and cross-correlation coefficient was calculated between them (I). See the Supplementary Figure 1 for the right hemisphere. MT: middle temporal area also known as V5; MST: medial superior temporal area; FST: fundal superior temporal area; LO1-3: area lateral occipital 1-3; PIT: posterior infero temporal; FFC: fusiform face complex; PeEc: perirhinal ectorhinal cortex; PHA: parahippocampal area; VMV: ventromedial visual area; VVC: ventral visual complex; POS1: parieto-occipital sulcus area 1; ProSt: prostriate area; A19DI: area 19 of cortex dorsointermediate part; A19M: area 19 of cortex medial part; Ent: entorhinal cortex; TLO: temporal area TL occipital part; TPO: temporo-parieto-occipital association area.

### Intra- and inter-species functional correspondence

To identify the intra- and inter-species functional correspondence in early and high-order visual areas, we created the volume-of-interests (VOIs) based on the inter-scan correlation maps (r > 0.2) using the multi-modal cortical parcellation atlas (Glasser et al., 2016) for humans and the Paxinos atlas for marmosets (Liu et al., 2021) (Fig. 1G and 1H; please see the Suppl. Fig. 1E and 1F for the right VOIs). The time courses were extracted from all VOIs and cross-correlation coefficients were calculated between all human and marmoset VOIs (see Fig. 1I and Methods section). The correlation matrices are shown in Suppl. Fig. 2. We found similar functional responses in human face-specific areas including the peri-entorhinal and ectorhinal cortex (PeEc), fusiform face complex (FFC), and posterior inferotemporal areas (PIT) (Augustinack et al., 2013; Gauthier et al., 2000; Glasser et al., 2016; Kanwisher et al., n.d.; Tsao et al., 2008) and in areas V4 transitional part (V4T), fundal superior temporal area (FST), and TE3 in marmosets. These marmoset areas are known to correspond to face patches based on task-based fMRI studies (Hung et al., 2015; Schaeffer et al., 2020b).

### Identification of marmoset areas corresponding to human face-, body-, and scene-specific areas

As a validation of our method, we next asked whether the previously described marmoset face patches (Hung et al., 2015; Schaeffer et al., 2020b) could be detected by comparing the time courses across the marmoset brain to the time course in one of the human face patches, PeEc (Glasser et al., 2016) also known as anterior face patch or Brodmann area 35/36 (Augustinack et al., 2013; Tsao et al., 2008). Calculating the cortex-wise correlation with the time course in the human face patch revealed discrete patches of the activity in both human (Red regions in Fig. 2A and 2C) and marmoset occipital and temporal areas (Red regions in Fig. 2B and 2D). There were four main clusters in temporal areas. Each cluster (F1, F2, F3, and F4) in humans consisted of PeEc and TF for F1, FFC, PH, and TE2 posterior area (TE2p) for F2, LO2, PIT, and V4T for F3, and middle temporal area (MT) and medical superior temporal area (MST) for F4 (Fig. 2C). Each cluster (F1, F2, F3, and F4) in marmosets consisted of A36 and entorhinal cortex (Ent) for F1, TE3 and temporo-parieto-occipital association area (TPO) for F2, FST and Pga-IPa for F3, and MT, MST, and V4T for F4 (Fig. 2C). The locations of these patches closely resembled those reported in the previous macaque (Hesse and Tsao, 2020; Landi and Freiwald, 2017; Tsao et al., 2008, 2003; Weiner and Grill-Spector, 2015) and marmoset fMRI studies (Hung et al., 2015; Schaeffer et al., 2020b), but we also found statistically significant activation in marmoset A36. This area has not been found in previous marmoset face fMRI studies, but it cytoarchitectonically corresponds to the anterior face patch in humans (Augustinack et al., 2013; Glasser et al., 2016; Rajimehr et al., 2009; Tsao et al., 2008). These marmoset areas were also found when we used the other human face patches, FFC and PIT as seed regions (Suppl. Fig. 5). This mapping of the face patches under more naturalistic viewing conditions demonstrates that it is possible to use this approach to identify marmoset areas that have the same function as specific human areas. All correlation coefficient maps with human VOIs are shown in Suppl. Figure 3–6, and the peak locations of these maps were summarized in Figure 3.

**Fig. 2.**
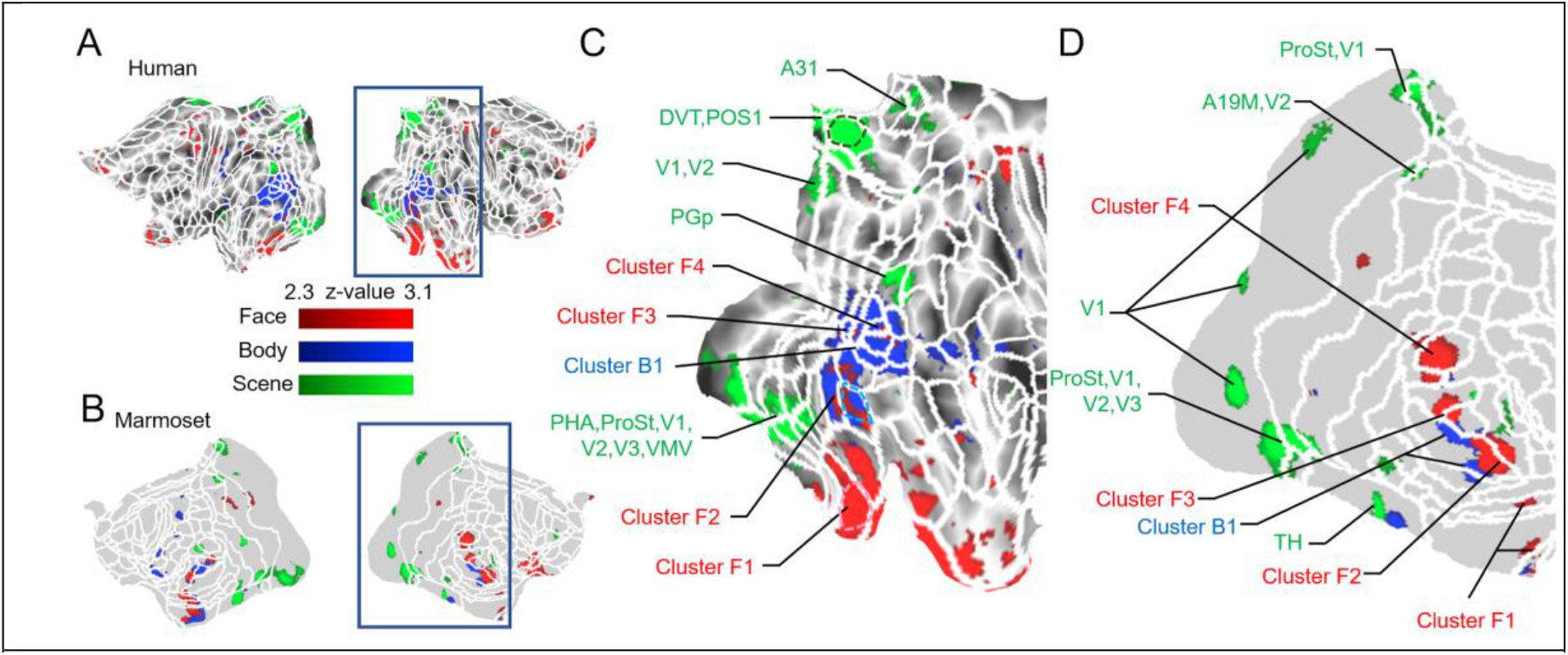
Correlation maps (z-score maps) in human (A) and marmoset (B) brains with human face-specific (PeEc indicated by pink dashed line in Fig. C), body-specific (TE2p indicated by light blue dashed line in Fig. C), and scene-specific (POS1 indicated by green dashed line in Fig. C) areas were presented on each flattened map. The figure C and D show the same data focusing around the occipital and temporal regions (areas surrounded by blue squares in Fig. A and B) in right hemispheres. White lines indicate the borders of the multi-modal cortical parcellation atlas (Glasser et al., 2016) for humans and Paxinos atlas for marmosets (Liu et al., 2021).

**Fig. 3.**
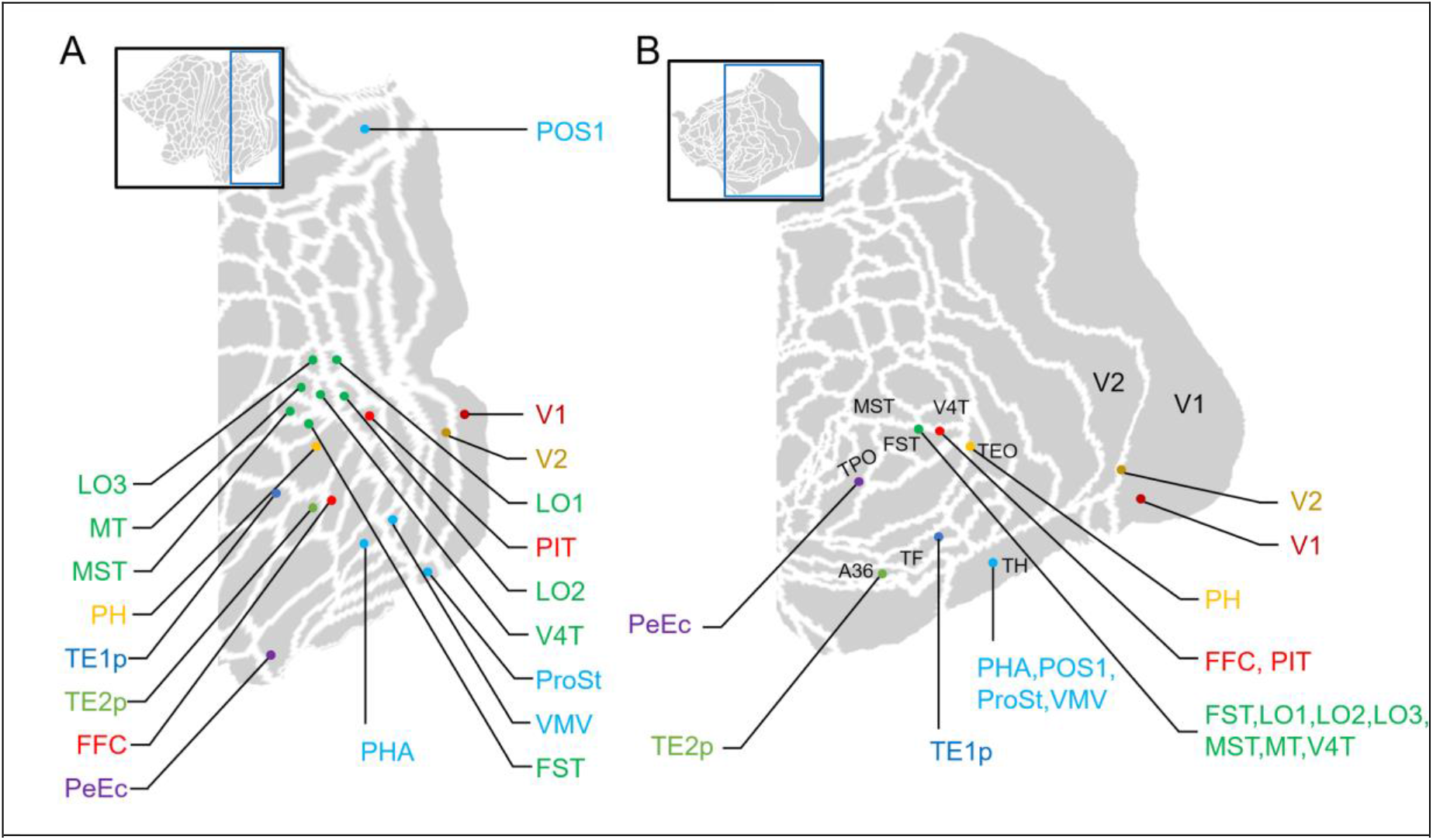
The peak locations of human (A) and marmoset (B) correlation maps with the time course in each human volume-of-interest (VOI) (presented by color labels). Note that seven out of 26 human VOIs (V3, V3A, V3B, V3CD, V4, V8, and VVC) were not presented due to the differences of the peak locations between left and right hemisphere. The maps were focused on the areas around the occipital regions (areas surrounded by blue squares in the top left panels) in left hemispheres. White lines indicate the borders of the multi-modal cortical parcellation atlas (Glasser et al., 2016) for humans and Paxinos atlas for marmosets (Liu et al., 2021).

Next, we sought to examine which marmoset areas correspond to body- and scene-specific areas in humans. To do so, we seeded the human TE2p (Glasser et al., 2016) and parieto-occipital sulcus area 1 (POS1: Glasser et al., 2016; Glasser and Van Essen, 2011), also known as retrosplenial cortex (Nasr et al., 2011) to identify marmoset body- and scene-specific areas, respectively. Body-specific areas have already been described in one previous task-based fMRI study in marmosets (Hung et al., 2015), and these areas are located adjacent to face patches, similar as in humans and macaque monkeys (Peelen and Downing, 2007; Pinsk et al., 2009; Tsao et al., 2003). Scene-specific areas have not yet been identified in marmosets.

The time course in the human body-specific area (TE2p) was highly correlated with parts of face specific areas (FFC and PIT) and the posterior superior temporal sulcus (pSTS) in humans, which is known as one of body patches (Pinsk et al., 2009) (Blue regions in Fig 2A and 2C). The time course in the human body patch was also highly correlated with the areas adjacent to the face patches in marmosets (Blue regions in Fig 2B and 2D).

The time course in the human scene-specific area (POS1) was highly correlated with A31, dorsal visual transitional area (DVT), PG posterior part (PGp), parahippocampal area (PHA), prostriate area (ProSt), ventromedial visual area (VMV), V1, V2 and V3 in humans (Green regions in Fig. 2A and 2B), and A19 medial (A19M), ProSt, TH (also known as PHA), V1, V2, V3 in marmosets (Green regions: Fig. 2C and 2D). This tendency was similar when we applied another human scene patches, PHA as a seed region (Suppl. Fig. 5). The locations of scene-specific areas in humans closely resembled those reported in previous human studies (Aguirre et al., n.d.; Epstein and Kanwisher, 1998; Ishai et al., 2000; Maguire, 2012). The activation areas in marmosets were broadly cytoarchitectonically consistent with human activation areas.

### Relative contribution of different features in face patches

To evaluate what features in the movie activate marmoset regions functionally corresponding to human face and body patches, we first identified for each TR interval (1.5 s) of the movie whether a marmoset, other animals (humans, owls, capybaras, dogs, cats, pigeons, roosters, frogs, scorpions, ants), or no animals were visible. These three pseudo-event related designs were convolved with a hemodynamic response function (HRF) using FSL FEAT (Fig. 4A). We then calculated the correlation coefficients between these predicted design and the time courses in each face and body patch. We found that the human face patches were more active when they watched not only marmosets, but also the other animals compared to no animals (Fig. 4B; p < 0.05 for cluster F1, p < 0.01 for cluster F2, F3, and F4; analysis of variance (ANOVA) with Bonferroni post-hoc correction). This tendency was also found for the human body patches (Fig. 4B; p < 0.01 for cluster B1; paired t-test with Bonferroni post-hoc correction). Unlike humans, marmoset face patches were more active when they watched marmosets (Fig. 4C; p < 0.05 for cluster F1 and F2, p < 0.01 for cluster F4; ANOVA with Bonferroni post-hoc correction) than when they watched the other animals. As in humans, the marmoset body patches were activated when they watched both marmosets and the other animals compared to no animals (Fig. 4B; p < 0.05; paired t-test with Bonferroni post-hoc correction).

**Fig. 4.**
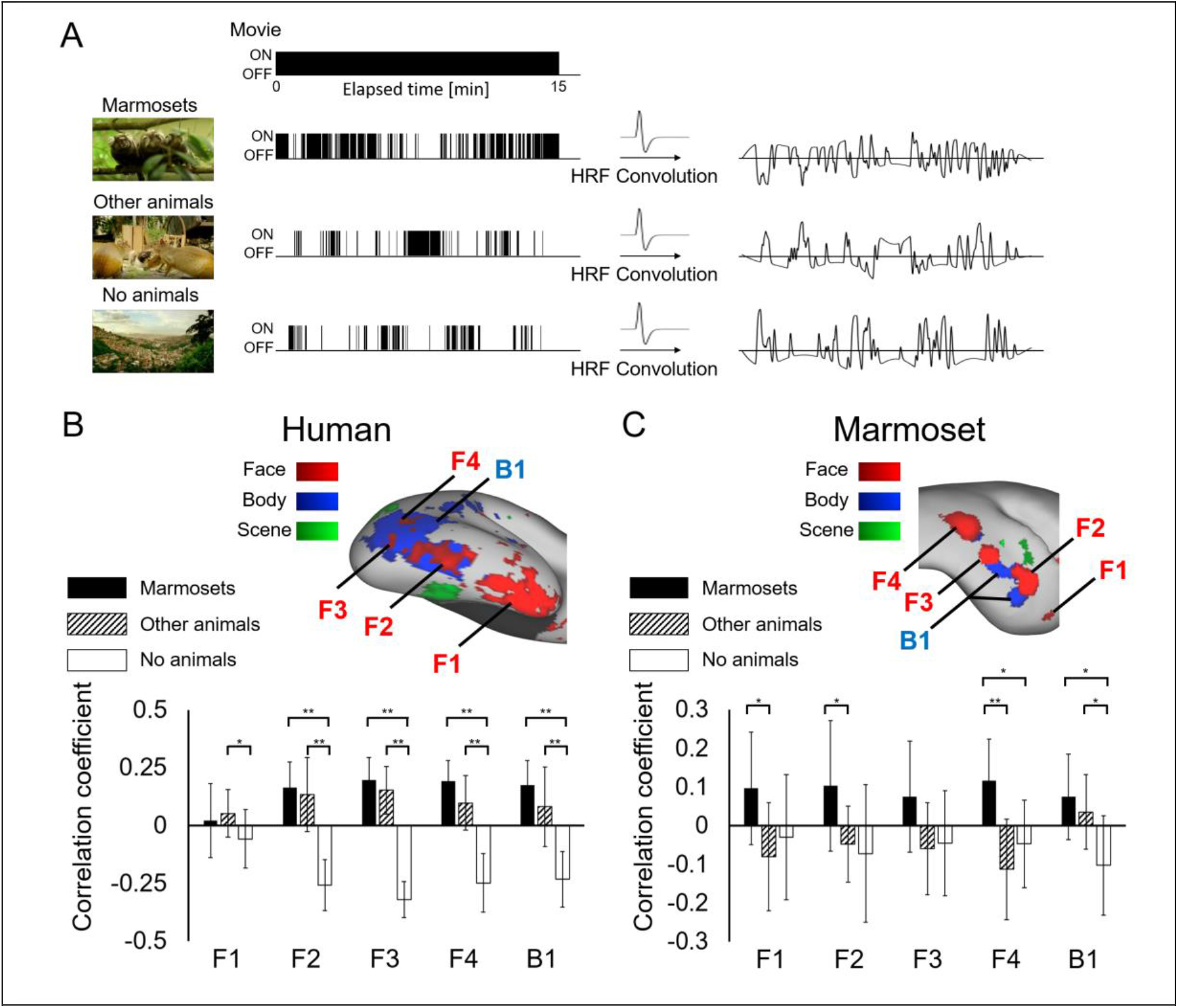
Correlations of the time course in each face or body patch with the pseudo-event related design. We first identified which animals (marmosets, other animals, or no animals) we can find in each TR (1.5 s) in the movie clip and were classified as 0 (OFF) and 1 (ON). These pseudo-event related designs were convolved by hemodynamic response function and were applied as regressors of interest (A). The locations of the clusters are shown on the cortical surfaces (F1 to F4 for face-specific clusters and B1 for body-specific cluster), which is the same data as shown in Fig. 2C and 2D. The black, slash, and white bars indicate the correlation coefficient values with the features of marmosets, other animals, and no animals, respectively. One and two asterisks indicate significant differences of p<0.05 and p<0.01 using analysis of variance (ANOVA) with Bonferroni post-hoc correction, respectively. Error bars indicate the standard deviations.

### Identification of human areas corresponding to marmoset parietal areas

As shown in Fig.1, both human and marmoset parietal areas also had high reproducibility between scans. We next asked which marmoset areas correspond to each human parietal area using the same method as above visual areas. However, for most human parietal areas we could not find correlations with parietal areas in marmosets (Suppl. Fig. 7-8). Therefore, we next identified human parietal areas that correlated with each marmoset area surrounding the intraparietal sulcus (IPS) (anterior intraparietal parietal area (AIP), lateral intraparietal parietal area (LIP), medial intraparietal area (MIP), PE, PG, occipito-parietal transitional area (OPt), ventral parietal area (VIP)). The analysis showed that the peak locations of the human correlation maps were located in area PG inferior part (PGi) for the marmoset anterior parietal regions (AIP, OPt, PE, and PG), and in the human area PG superior part (PGs) for the marmoset parietal regions LIP, MIP, and VIP (Fig. 5).

**Fig. 5.**
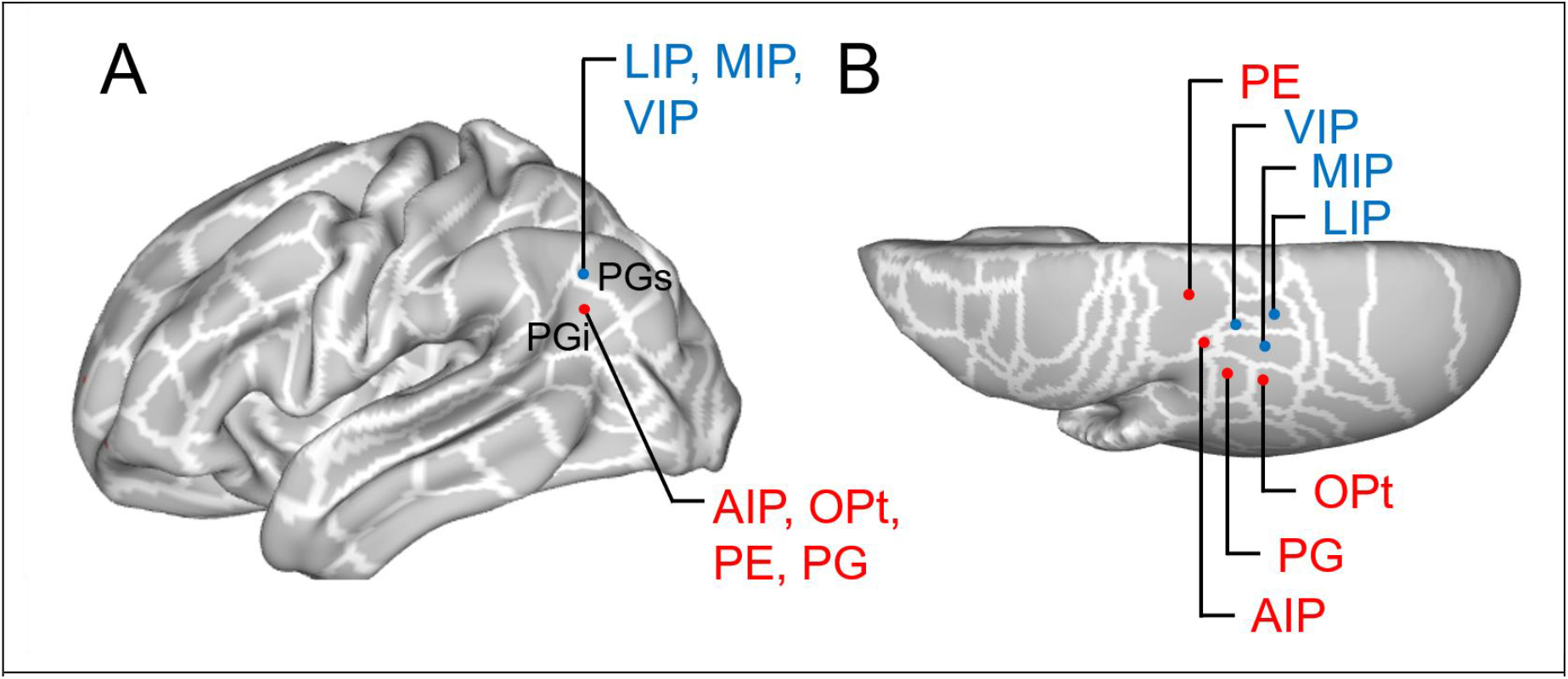
The peak locations of human (A) and marmoset (B) correlation maps with the time course in each marmoset parietal volume-of-interest (VOI) (presented by red or blue labels). White lines indicate the borders of the multi-modal cortical parcellation atlas (Glasser et al., 2016) for humans and Paxinos atlas for marmosets (Liu et al., 2021) AIP: anterior intraparietal area; LIP: lateral intraparietal area; MIP: medial intraparietal area; OPt: occipito-parietal transitional area; VIP: ventral intraparietal area

## Discussion

### Possible homology between human and marmoset face areas

In this study, we aimed to identify areas in the marmoset that functionally correspond to high-order visual areas in humans, in particular face-, body-, and scene-specific areas. To do so, we presented human participants and marmoset monkeys with a naturalistic movie during fMRI acquisition at ultra-high field. We found functional responses in human face patches (i.e. PeEc, FFC, and PIT) that were in good agreement with those in marmoset dorsal face patches (Hung et al., 2015; Schaeffer et al., 2020b) (i.e. an anterior dorsal (AD), a middle dorsal (MD), and a posterior dorsal (PD) areas). In addition, we observed a functional correspondence between the PeEc in humans and area 36 in marmosets, which had not yet been described in previous studies. The human PeEc in the multi-modal parcellation atlas (Glasser et al., 2016) corresponds to the AFP and Brodmann area 36 (Augustinack et al., 2013; Glasser et al., 2016; Rajimehr et al., 2009; Tsao et al., 2008). Thus, area 36 in marmosets seems to correspond to the AFP in humans. To further explore the functions of marmoset face patches, we also identified what features in the movie activated the different patches. These analyses demonstrated that face patches in marmosets responded to marmosets but not other animals, whereas human face patches exhibited similar responses to marmosets and other animals. This tendency was similar to a previous NHP study in macaques that showed that macaque face patches responded more than twice as strongly to macaque faces compared to human faces, whereas human face patches responded similarly to the presentation of both human and macaque faces (Tsao et al., 2003). This indicates that both macaque and marmoset face patches preferentially respond to con-specific faces.

Previous systematic mapping of human and macaque visual cortices onto each other have revealed an overall shift of areas ventrally from the superior temporal sulcus (STS) in humans compared to macaque monkeys (Orban et al., 2004), corresponding to an overall areal expansion in this region (Van Essen and Dierker, 2007). Along with this shift of face areas, macaque face patches are located more dorsally compared to human face patches (Tsao et al., 2008). As in macaque monkeys, marmoset face patches are also located dorsally compared to human face patches (Hung et al., 2015; Schaeffer et al., 2020b). Given the locations of marmoset face patches relative to each other, the marmoset AD and MD face patches along the marmoset STS (corresponding to the clusters F2 and F3 in Fig. 2D) are in a similar position as the macaque anterior fundus (AF) and middle fundus (MF) face patches, respectively (Tsao et al., 2008). Their positions also bear resemblance to two face patches in anterior and posterior STS in humans (Carlin et al., 2011; Pitcher et al., 2011), although the correspondence between the human and non-human primate is still in a matter of speculation (Yovel and Freiwald, 2013). The marmoset PD patch (corresponding to the cluster F4 in Fig. 2D), which is located in MT, MST, and V4T, is in a similar position as PL in macaque V4T (Janssens et al., 2014), and an occipital facial area (OFA) in human PIT (Abdollahi et al., 2014; Tsao et al., 2008). On the other hand, we could not detect ventral face patches in marmosets (i.e. posterior ventral (PV) and middle ventral (MV) (Hung et al., 2015; Schaeffer et al., 2020b)). The face patches along with the dorsoventral axis show different selectivity to natural motion (Fisher and Freiwald, 2015; Zhang et al., 2020). Ventral face patches have a preference for rapidly varying face stimuli, whereas dorsal face patches have a preference for natural motion. These differences in selectivity may present the detection of ventral face patches with the movie employed here.

### Body and scene patch systems in humans and marmosets

The functional responses in the human body-specific patch (TE2p) were in good agreement with those in the marmoset area located between the face patches AD and MD. This area is already known as a marmoset body-specific patch based on a previous electrocorticography and fMRI study (Hung et al., 2015). In addition, we also found that the human TE2p correlated with the marmoset area adjacent to the anterior part of AD, which had not been found yet. This finding is broadly consistent with previous findings in macaques and humans of body-selective areas located adjacent to face patches (Downing, 2001; Peelen et al., 2007; Peelen and Downing, 2007; Tsao et al., 2003; Weiner and Grill-Spector, 2015).

In humans, previous functional MRI studies (Downing, 2001; Peelen et al., 2007; Tsao et al., 2003; Weiner and Grill-Spector, 2013) have described three visual cortical regions that are more active during the presentation of scenes or isolated houses compared with the presentation of other visual stimuli such as faces, objects, body parts, or scrambled scenes. Typically, these human brain regions are located in the parieto-occipital sulcus area 1 (POS1), parahippocampal area (PHA), and transverse occipital sulcus (TOS). In this study, we found that functional responses in human POS1 and PHA corresponded to those in marmoset retrosplenial cortex (A19M) and in the parahippocampal area (TH), suggesting a homologous neural architecture for scene-selective regions in visual cortex for humans and marmosets.

### Functional correspondence in parietal and motion-selective visual areas

As described above, we identified areas in the marmoset that functionally corresponded to face-, body-, and scene-specific areas in humans. Next, we sought to examine which marmoset areas show functional correspondences to other visual areas in humans. Our results demonstrate that area MT and its surrounding areas in humans (FST, LO1, LO2, LO3, MST, and V4T) corresponded to a region at the border between FST, MST, and V4T. Area MT in both New-World and Old-World monkeys has direction-selective neurons (Born and Bradley, 2005), and are considered to play an important role in the motion perception, direction selectivity, and speed turning (DeYoe and Van Essen, 1988; Lui and Rosa, 2015; Saito et al., 1986). The major output of MT is in the cortical areas surrounding it (i.e. FST, MST, and V4T) ((Ungerleider and Desimone, 1986) for macaques; (Abe et al., 2018) for marmosets). Therefore, the movie may not be suitable to identify small functional differences between these regions. The movie we used was selected to identify face-, body, and scene patches. To better identify motion-related areas, it would be ideal to use a movie in which stimuli move in various directions, speeds, and rotations but this is beyond the scope of this study.

Although we could not identify functional differences between most marmoset parietal areas, the anterior (AIP, OPt, PE, and PG) and posterior (LIP, MIP, and VIP) parietal regions in marmosets were correlated with PGi and PGs, respectively. As described in numerous studies, human PG is a part of the default mode network (DMN), while the areas surrounding the IPS (e.g. area LIP) is a part of the attention network (ATN) (Glasser et al., 2016; Ji et al., 2019; Thomas Yeo et al., 2011). In addition, the time courses of blood-oxygen-level-dependent (BOLD) signals in human PG are anti-correlated with those in task positive regions (e.g. frontal eye field, LIP) even under resting-state (Fox et al., 2005). In marmosets, one of the core regions of the DMN is the cortex surrounding the IPS (Belcher et al., 2013; Hori et al., 2020b; Liu et al., 2019). A functional imaging study in marmosets showed that these areas were deactivated during a visual stimulation task relative to the period when a black screen was presented (Liu et al., 2019). On the other hand, the area surrounding the marmoset IPS have also been linked to the attention-like frontoparietal network by several RS-fMRI studies (Ghahremani et al., 2016; Hori et al., 2020b). It is also known that electrical microstimulation in the areas surrounding the IPS evoke saccadic eye movements (Ghahremani et al., 2019) and that single neurons in the region show neural correlates for the gap effect in saccadic eye movement tasks (Ma et al., 2020). As such, it seems that parietal areas surrounding the IPS in the marmoset are involved in both default mode and attention networks. Taken together with our finding that the marmoset parietal area corresponded to the parietal regions in the human DMN, the DMN might be predominantly engaged during movie viewing in the marmoset. Indeed, our results showed that the human PG was correlated with the posterior cingulate cortex (PCC), dorso-lateral prefrontal cortex (dlPFC), and parts of anterior cingulate cortex (ACC) and temporal areas (Suppl. Fig. 9), and these areas overlap with the human DMN (Glasser et al., 2016; Ji et al., 2019; Thomas Yeo et al., 2011) which is deactivated during attention demanding tasks (Fox et al., 2005; Sani et al., 2021).

### Differences between human and marmoset frontal areas

We also observed consistent activation in marmoset frontal areas, in particular, the medial PFC (mPFC; A25, A32), ventro-lateral PFC (vlPFC; A45, A47), and dorso-medial PFC (dmPFC; A8b, A9) (Fig. 1D-F). This distribution closely resembled a recently-identified network for social interaction processing (Sliwa and Freiwald, 2017). The authors found that the macaque mPFC (A10mr, A24b, A32), vlPFC (A12, A44, A47, OPro), and dmPFC (A8b, A9m, F6) are more activated when the monkey watched a social video (in which two monkeys were interacting with each other) compared to a non-social video (in which two monkeys were separately acting in their own environment). As such, consistent activation in the marmoset frontal areas might reflect social interaction processing during movie viewing. Although humans possess a similar social interaction network (Mars et al., 2012; Schilbach et al., 2008), the social interactions of marmosets might not have been enough to consistently activate this network in our human participants.

### Conclusion

In summary, we identified marmoset areas that functionally corresponded to human face-, body-, and scene-processing areas using MD-fMRI. The locations of these marmoset areas relative to each other were broadly consistent with those of human areas, suggesting that high-order visual processing might be a conserved feature between humans and New World marmoset monkeys. These findings further strengthen the marmoset as a powerful non-human primate model for visual and social neuroscience.

## Methods

### Subjects

Thirteen healthy volunteers (nine males and four females, 22-56 years) and four common marmosets (Three males and one female, 30-38 months,310-410 g) participated in this study. All surgical and experimental procedures for marmosets were in accordance with the Canadian Council of Animal Care policy and a protocol approved by the Animal Care Committee of the University of Western Ontario Council on Animal Care. All animal experiments complied with the Animal Research: Reporting *In Vivo* Experiments (ARRIVE) guidelines. Human volunteers were informed about the experimental procedures and provided informed written consent. This study was approved by the Ethics Committee of the University of Western Ontario.

### Animal preparation

All marmosets underwent a surgery to implant a head chamber to immobilize the head during MRI acquisition as described in previous reports (Johnston et al., 2018; Schaeffer et al., 2019c). Briefly, the marmoset was placed in a stereotactic frame (Narishige Model SR-6C-HT), and several coats of adhesive resin (All-bond Universal Bisco, Schaumburg, Illinois, USA) were applied using a microbrush, air dried, and cured with an ultraviolet dental curing light. Then, a dental cement (C&B Cement, Bisco, Schaumburg, Illinois, USA) was applied to the skull and to the bottom of the chamber, which was then lowered onto the skull via a stereotactic manipulator to ensure correct location and orientation. Each marmoset was trained over the course of three weeks to acclimatize to the MRI scanning environment and head-fixation system in a mock MRI environment following the procedures described by Silva et al. (Silva et al., 2011).

### Experimental setup

Both humans and marmosets watched a 15 min of naturalistic movie with no sound. The movie was edited from the BBC episode “Urban jungles” from the “Hidden Kingdoms” series. The episode was condensed to 15 min by removing the parts that were not related to marmosets living in Rio de Janeiro and removing some scenes that seemed to be less interesting to marmosets based on some pilot data (essentially scenes that did not show any marmosets for a while).

Human volunteers lay in a supine position and watched the movie presented via a rear projection system (Avotech SV-6011, Avotec Incorporated, Stuart, Florida, USA) through a surface mirror affixed to head coil (6.9° field of view-of-view from the center to the side of the screen). Each marmoset was fixed to the animal holder using a neck plate and a tail plate. The marmoset was then head-fixed in a sphinx position using fixation pins in the MRI room to minimize the time in which the awake animal was head fixed (Schaeffer et al., 2019). Once fixed, a lubricating gel (MUKO SM321N, Canadian Custom Packaging Company, Toronto, Ontario, Canada) was squeezed into the chamber and applied to the brow ridge to reduce magnetic susceptibility. For the marmosets, the movie was presented via a projector (Model VLP-FE40, Sony Corporation, Tokyo, Japan), reflected off of a first surface mirror, and back-projected onto a plastic screen that was velcroed to the front of the scanner bore (6.9° field of view-of-view from the center to the side of the screen). Both humans and marmosets freely watched the movie without any fixation constraints. In marmosets, the eyes were monitored using an MRI compatible camera (Model 12M-i, MRC Systems GmbH, Heidelberg, Germany) to make sure that the animals stayed awake. For both species, the movie was presented via Powerpoint on a MacBook Pro.

### Image acquisition

All imaging was performed at the Centre for Functional and Metabolic Mapping at the University of Western Ontario. Human images were acquired using a 68 cm head-only 7 T MRI scanner (Siemens Magnetom 7T MRI Plus, Erlangen, Germany) with an AC-84 Mark II gradient coil, an in-house 8-channel parallel transmit, and 32-channel receive coil (Gilbert et al., 2015). Functional images were acquired with a single run for each volunteer (624 volumes for each run), using gradient-echo based single-shot EPI sequence with the following parameters: TR = 1500 ms, TE = 20 ms, flip angle = 30°, field of view (FOV) = 208 × 208 mm, matrix size 104 × 104, voxel size 2.0 mm isotropic, slices = 60, GRAPPA acceleration factor (anterior-posterior) = 3, multiband factor = 2. For the first and last 18 sec (12 volumes for each), a fixation point was presented to the subject. Field map images were collected for EPI distortion correction with the following parameters: TR = 475 ms, TE1 / TE2 = 4.08 / 5.1 ms, flip angle = 35°, field of view (FOV) = 210 × 210 mm. A MP2RAGE structural image was also acquired for each subject with the following parameters: TR = 6000 ms, TE = 2.13 ms, TI1 / TI2 = 800 / 2700 ms, FOV = 240 × 240 mm, matrix size = 320 × 320, voxel size = 0.75 × 0.75 × 0.75 mm, slices per slab= 208, GRAPPA acceleration factor (anterior-posterior) = 3.

Marmoset images were acquired using a 9.4 T 31 cm horizontal bore magnet (Varian/Agilent, Yarnton, UK) and Bruker BioSpec Avance III HD console with the software package Paravision-6 (Bruker BioSpin Corp, Billerica, MA), a custom-built high-performance 15-cm-diameter gradient coil with 400-mT/m maximum gradient strength (Handler et al., 2020), and the 5-channel receive coil (Schaeffer et al., 2019c). Radiofrequency transmission was accomplished with a quadrature birdcage coil (12-cm inner diameter) built in-house. Functional images were acquired with 3-4 functional runs for each animal under the awake condition (624 volumes for each run), using gradient-echo based single-shot echo-planar imaging (EPI) sequence with the following parameters: TR = 1500 ms, TE = 15 ms, flip angle = 40°, field of view (FOV) = 64 × 64 mm, matrix size 128 × 128, voxel size 0.5 mm isotropic, slices = 42, bandwidth = 500 kHz, generalized autocalibrating parallel acquisition (GRAPPA) acceleration factor (anterior-posterior) = 2. As in humans, for the first and last 18 sec, a fixation point was presented, and the movie was presented for 15 min between those (600 TRs). A T2-weighted image (T2w) was also acquired for each animal using rapid imaging with refocused echoes (RARE) sequences with the following parameters: TR = 5500 ms, TE = 53 ms, FOV = 51.2 × 51.2 mm, matrix size = 384 × 384, voxel size = 0.133 × 0.133 × 0.5 mm, slices = 42, bandwidth = 50 kHz, GRAPPA acceleration factor (anterior-posterior) = 2.

### Image preprocessing

Data was preprocessed using FSL software (Smith et al., 2004). Raw MRI images were first converted to Neuro Informatics Technology Initiative (NIfTI) format (Li et al., 2016), and brain masks were created using the population-based brain template (MNI152 template for humans; MBM template for marmosets (Liu et al., 2021)). The brain-skull boundary was roughly identified from individual anatomical images using the brain extraction tool (FSL; BET) (Smith, 2002). Then, the brain template was linearly and non-linearly registered to the individual brain image using FMRIB’s linear registration tool (FLIRT) and FMRIB’s nonlinear registration tool (FNIRT) to more accurately create the brain mask. fMRI images were corrected for motion (FSL; FLIRT) and B0 geometric distortions (FSL; FEAT; only for humans). All fMRI images were finally normalized to the template using fMRI-to-structure and structure-to-template transformation matrices obtained by FLIRT and FNIRT, followed by spatial smoothing (3.0 mm for humans and 1.5 mm for marmosets) and temporal filtering (0.01-0.1 Hz).

### Inter-scan variability

To estimate the inter-scan variability of stimulus-driven activity, we calculated the voxel-by-voxel correlation across scans within each species. We first randomly separated the subjects into two groups and averaged the time courses voxel-by-voxel in each group. We then calculated the cross-correlation coefficient maps between two groups using Matlab (xcorr; The Math Works, Natick, MA). These processes were repeated 1000 times, and we obtained the average correlation coefficient maps. These maps were presented on the flattened cortical surface (Glasser et al., 2016, 2013) using the Connectome Workbench software (Marcus et al., 2011).

### Intra- and Inter-species activity correlations

To evaluate the intra- and inter-species activity correlations, the preprocessed fMRI datasets were first averaged across scans, and the cerebral cortical grey matter voxels were mapped to the surface (Glasser et al., 2016, 2013) using the Connectome Workbench software (Marcus et al., 2011). Then, the volume-of-interests (VOIs) in visual and parietal areas were manually created based on the multi-modal cortical parcellation atlas (Glasser et al., 2016) for humans and Paxinos atlas for marmosets (Liu et al., 2021) to avoid including the low reproducibility regions among scans (r < 0.2) based on the inter-scan variability maps (Fig. 1A and 1B). Finally, we obtained 26 and 25 VOIs for humans and marmosets, respectively (Fig. 1C, 1D for the left hemisphere and Suppl. Fig. 1 for right hemisphere). The time courses were extracted from each VOI, and intra- and inter-species correlation matrices were created by cross-correlating the time courses among two regions using Matlab (The Math Works, Natick, MA) (Fig. 1E).

### Identification of functional correspondent regions between humans and marmosets

To identify marmoset areas that have similar functional activations as human visual-related regions, we extracted the human time course in each 26 VOI and we created the correlation coefficient maps with these time courses. These correlation coefficient maps were Fisher Z-transformed for each of two species and were threshold at z = 2.3.

### Relative contribution of different features in face patches

To evaluate the functional similarities or dissimilarities of face and body-specific patches in humans and marmosets, we identified what features in the movie activated each region. To do so, we first visually identified which animals were present in each TR (1.5 s) in the movie (Fig. 3). These pseudo-event related designs were convolved with a canonical hemodynamic response function (HRF) using FSL FEAT. We then calculated the correlation coefficients between these predicted designs and the time courses in the face and body patches identified by correlations with human PeEc and TE2p in each subject. PeEc and TE2p are known as face-specific (Augustinack et al., 2013; Tsao et al., 2008) and body-specific areas in humans (Glasser et al., 2016; Peelen and Downing, 2007; Pinsk et al., 2009), respectively. The correlation coefficient values were finally averaged across subjects. The statistical differences were assessed using analysis of variance (ANOVA) with Bonferroni post-hoc correction.

## Acknowledgements

This work was supported by the Canadian Institutes of Health Research (FRN 148365, FRN 353372) and the Canada First Research Excellence Fund to BrainsCAN. We also thank Miranda Bellyou, Cheryl Vander Tuin and Hannah Pettypiece for animal preparation and care and Dr. Alex Li and Trevor Szekeres for scanning assistance.

## Supplementary figures

**Suppl. Fig. 1.**
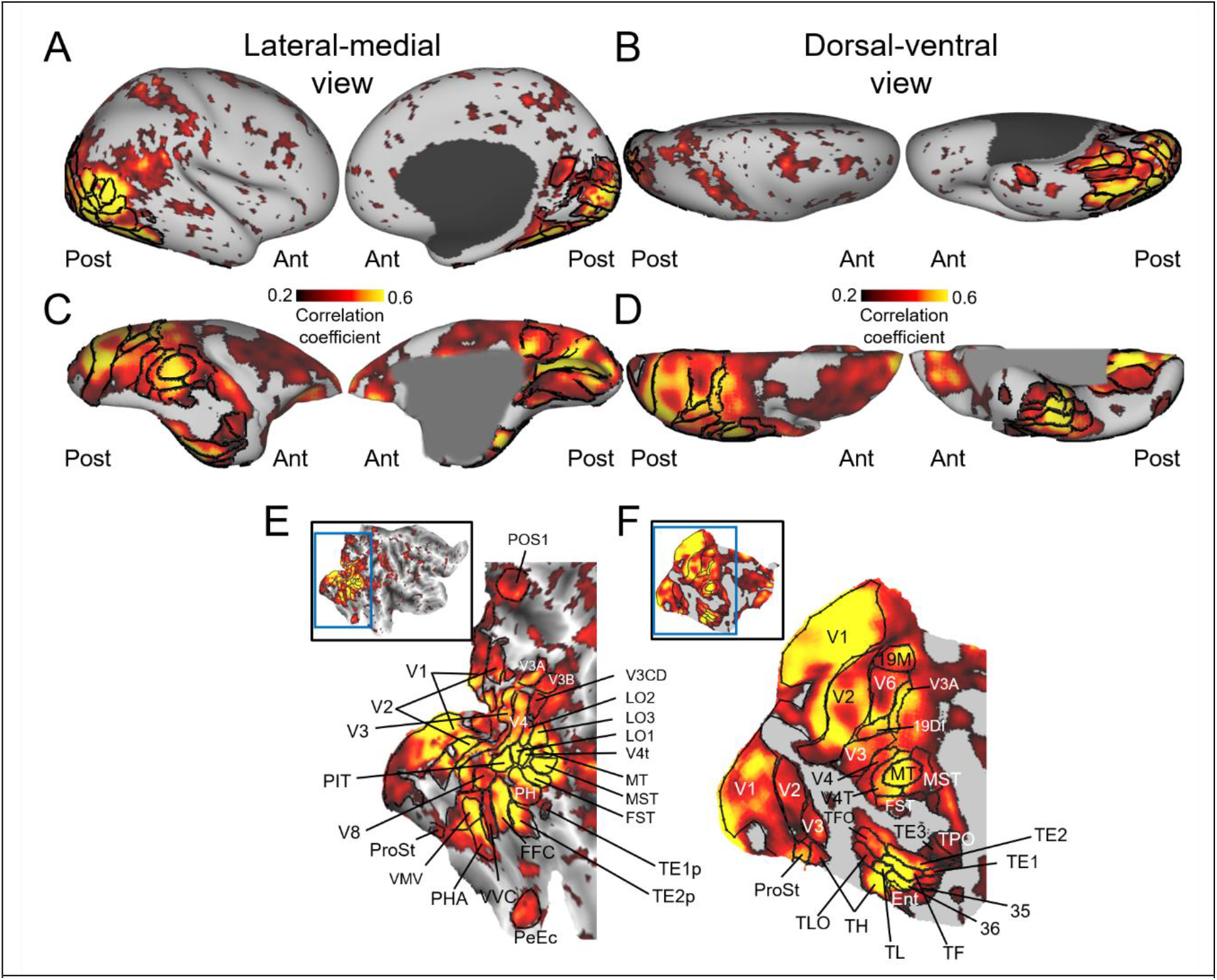
Inter-scan correlation map of brain activity during movie viewing. Spatial maps of correlated brain activity across 13 scans (13 subjects) (A) and across 14 scans (4 subjects), mapped on right cortical surface (lateral-medial view (A, C) and dorsal-ventral view (B, D)). The volume-of-interests (VOIs) in visual-related areas were manually created based on the multi-modal cortical parcellation atlas (Glasser et al., 2016) for humans and Paxinos atlas for marmosets (Liu et al., 2021), so as not to include the low correlation areas among scans (r < 0.2) (E for humans, and F for marmosets). MT: middle temporal area also known as V5; MST: medial superior temporal area; FST: fundal superior temporal area; LO1-3: area lateral occipital 1-3; PIT: posterior infero temporal; FFC: fusiform face complex; PeEc: perirhinal ectorhinal cortex; PHA: parahippocampal area; VMV: ventromedial visual area; VVC: ventral visual complex; POS1: parieto-occipital sulcus area 1; ProSt: prostriate area; A19DI: area 19 of cortex dorsointermediate part; A19M: area 19 of cortex medial part; Ent: entorhinal cortex; TLO: temporal area TL occipital part; TPO: temporo-parieto-occipital association area.

**Suppl. Fig. 2.**
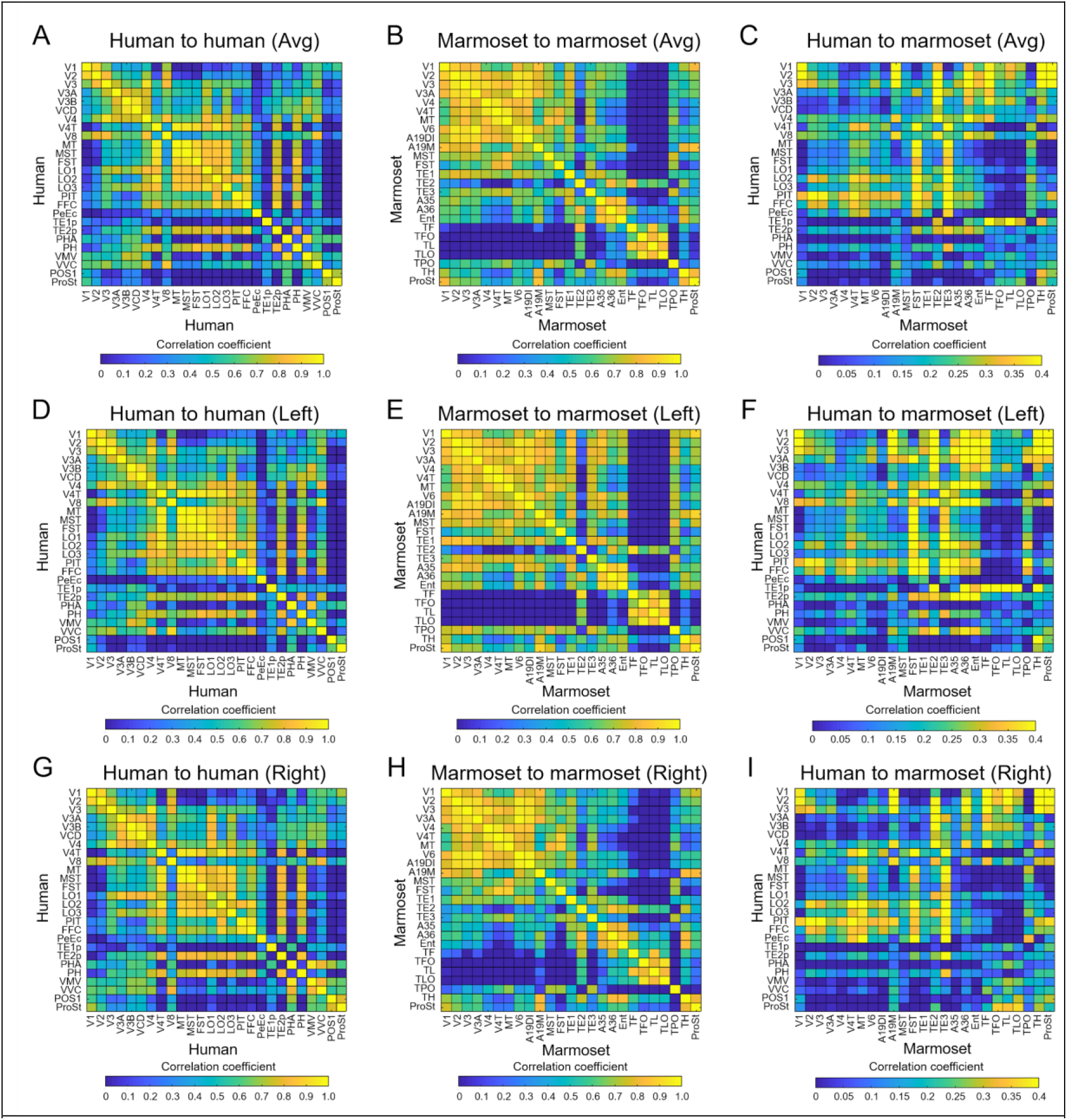
Intra-species correlation matrices for humans (A, D, G) and for marmosets (B, E, H) were shown in a range from 0 to 1 and inter-species correlation matrix (C, F, I) was shown in a range from 0 to 0.4. Top figures (A, B, C) were calculated by averaging the matrices of left (D, E, F) and right hemispheres (G, H, I), respectively.

**Suppl. Fig. 3.**
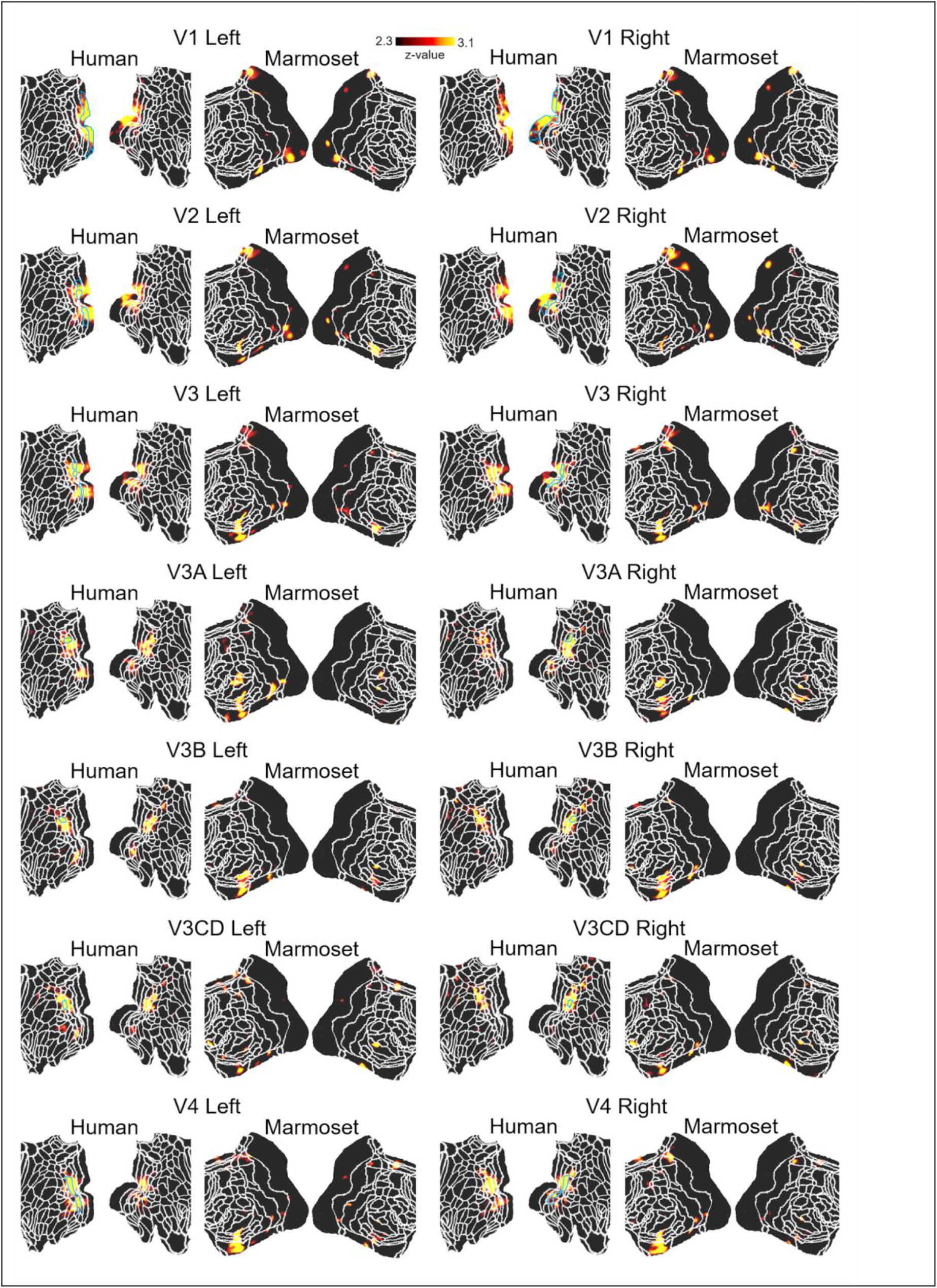
Correlation maps (z-score maps) in human and marmoset brains from each human volume-of-interest (presented above the map). The maps from fourteen of 52 VOIs (other 38 in Suppl. figs. 4–6) were shown as thresholded z-score maps on each flattened cortex. The area surrounded by sky blue line indicates the seed regions. White lines indicate the borders of the multi-modal cortical parcellation atlas (Glasser et al., 2016) for humans and Paxinos atlas for marmosets (Liu et al., 2021).

**Suppl. Fig. 4.**
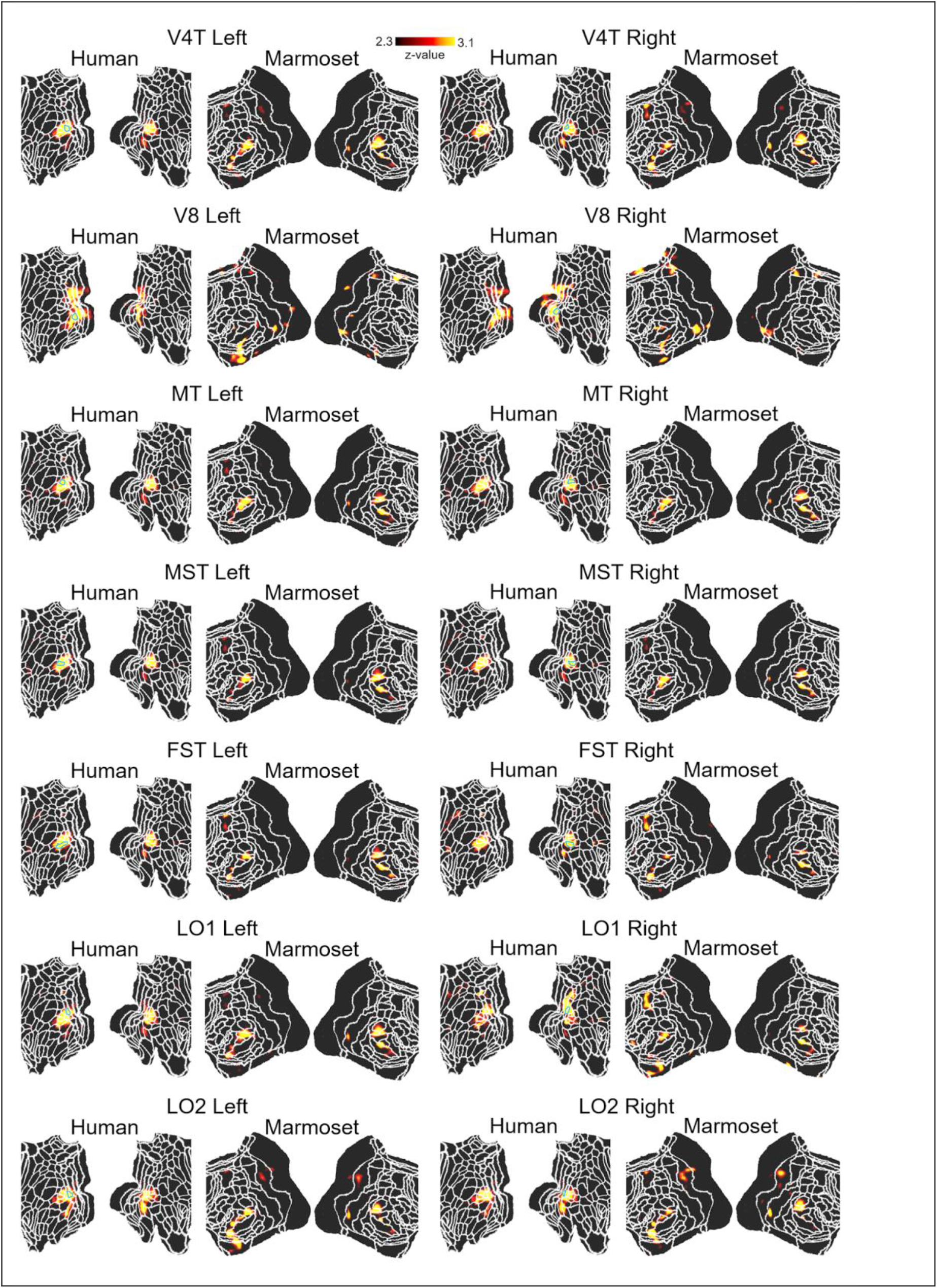
Correlation maps (z-score maps) in human and marmoset brains from each human volume-of-interest (presented above the map). The maps from fourteen of 52 VOIs (other 38 in Suppl. figs. 3, 5, and 6) were shown as thresholded z-score maps on each flattened cortex. The area surrounded by sky blue line indicates the seed regions. White lines indicate the borders of the multi-modal cortical parcellation atlas (Glasser et al., 2016) for humans and Paxinos atlas for marmosets (Liu et al., 2021).

**Suppl. Fig. 5.**
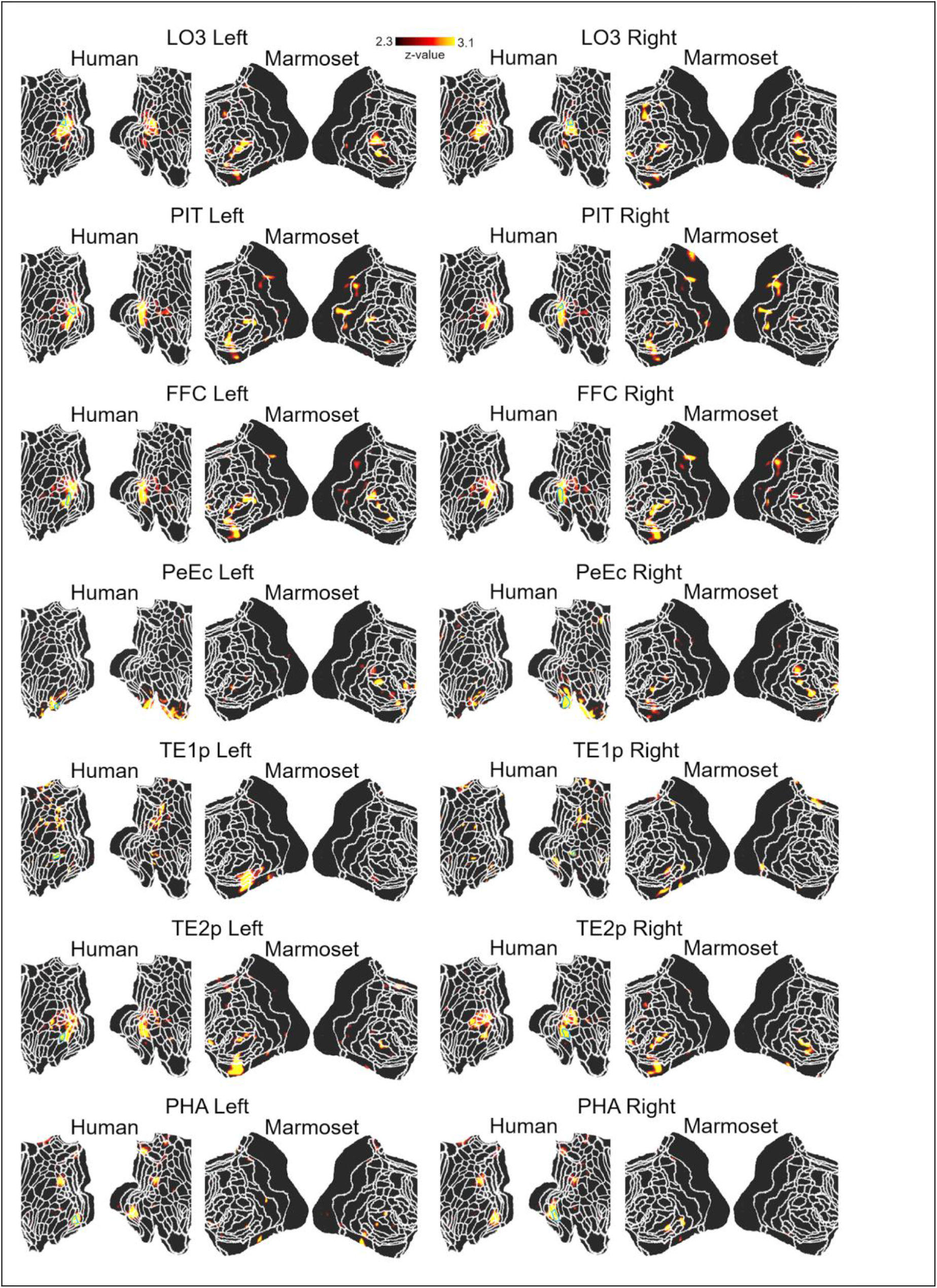
Correlation maps (z-score maps) in human and marmoset brains from each human volume-of-interest (presented above the map). The maps from fourteen of 52 VOIs (other 38 in Suppl. figs. 3, 4, and 6) were shown as thresholded z-score maps on each flattened cortex. The area surrounded by sky blue line indicates the seed regions. White lines indicate the borders of the multi-modal cortical parcellation atlas (Glasser et al., 2016) for humans and Paxinos atlas for marmosets (Liu et al., 2021).

**Suppl. Fig. 6.**
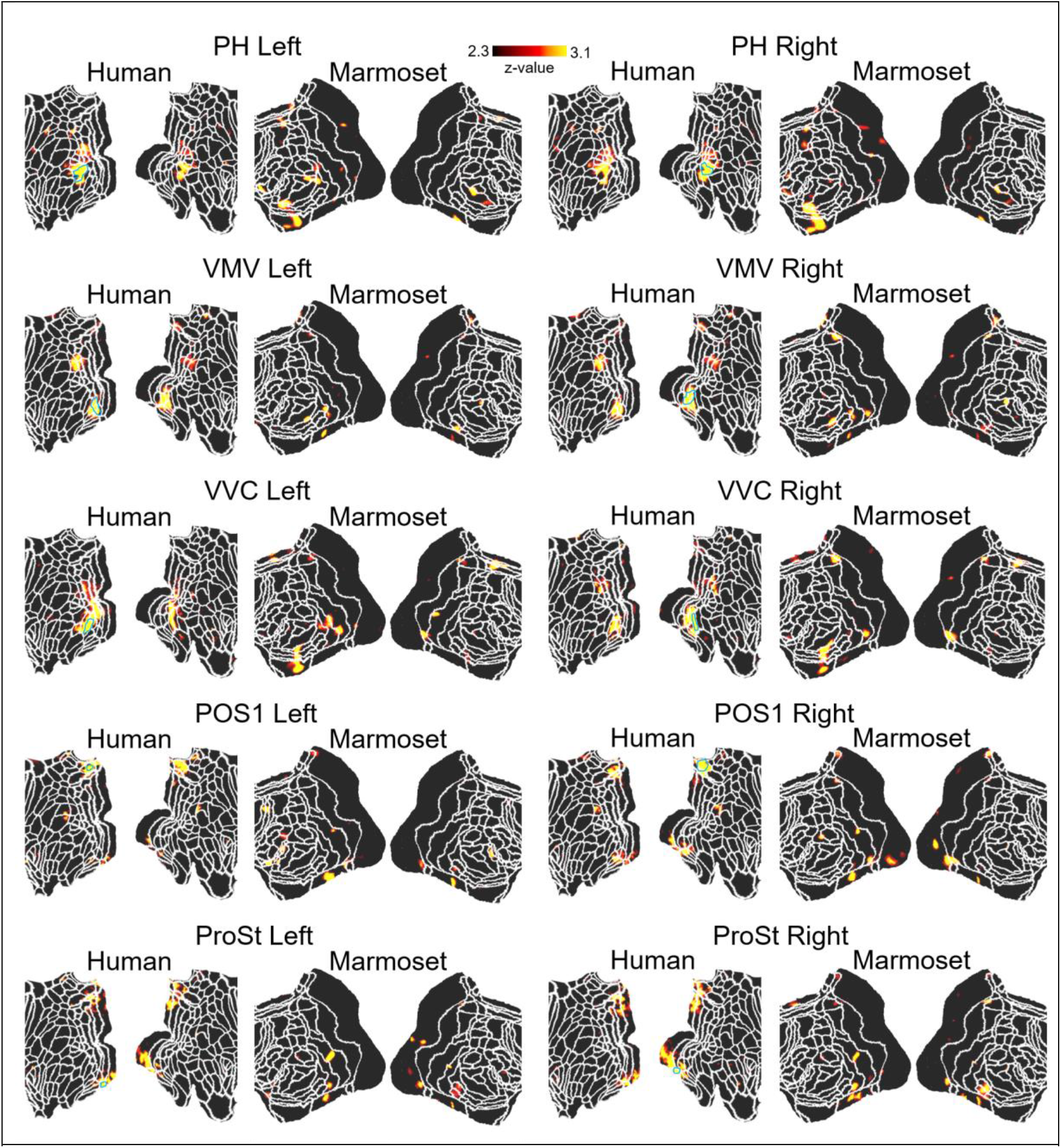
Correlation maps (z-score maps) in human and marmoset brains from each human volume-of-interest (presented above the map). The maps from ten of 52 VOIs (other 42 in Suppl. figs. 3–5) were shown as thresholded z-score maps on each flattened cortex. The area surrounded by sky blue line indicates the seed regions. White lines indicate the borders of the multi-modal cortical parcellation atlas (Glasser et al., 2016) for humans and Paxinos atlas for marmosets (Liu et al., 2021).

**Suppl. Fig. 7.**
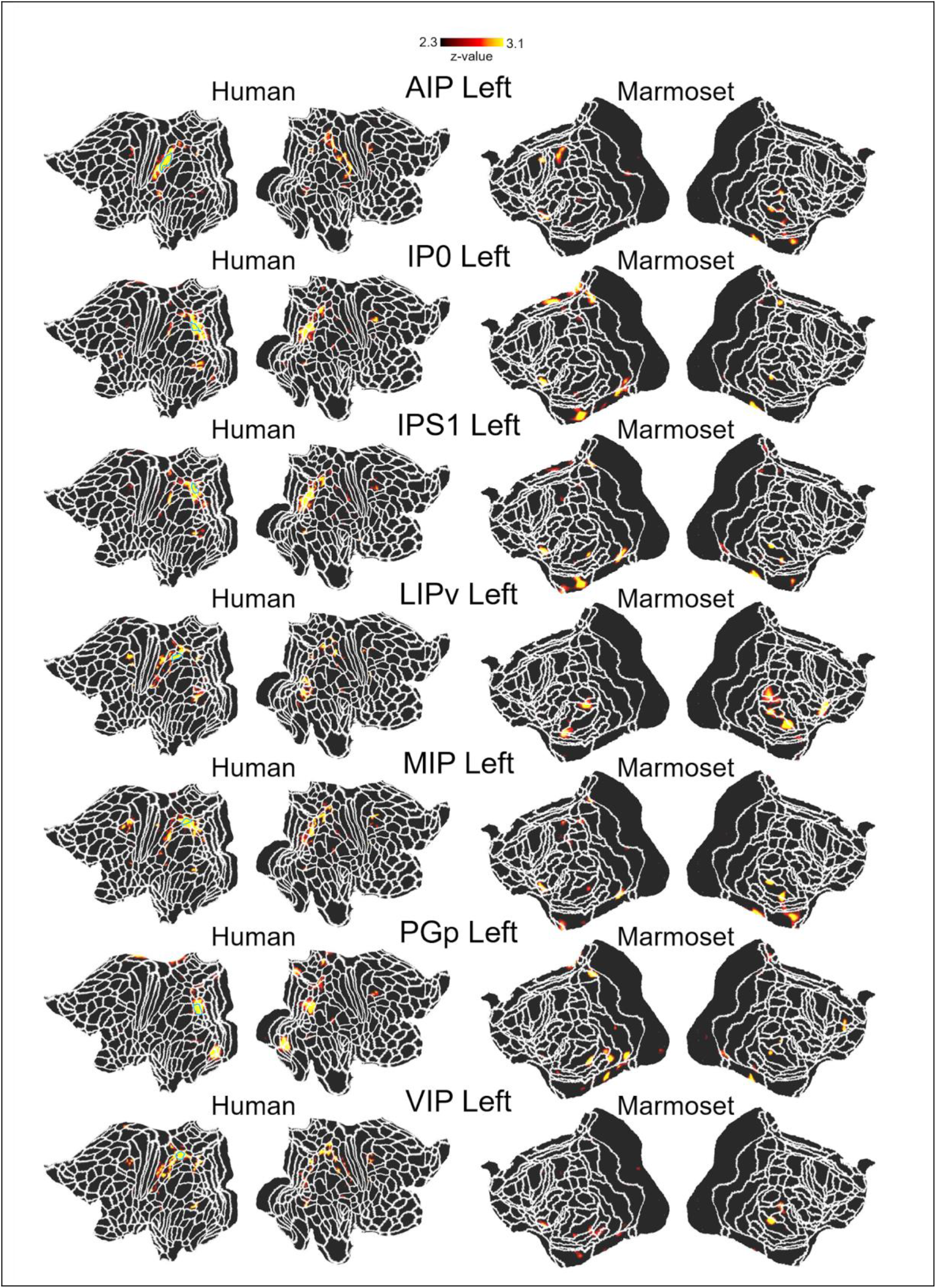
Correlation maps (z-score maps) in human and marmoset brains from each human left parietal volume-of-interest (presented above the map). The maps were shown as thresholded z-score maps on each flattened cortex. The area surrounded by sky blue line indicates the seed regions. The maps for the right VOIs were shown in Suppl. Fig. 8. White lines indicate the borders of the multi-modal cortical parcellation atlas (Glasser et al., 2016) for humans and Paxinos atlas for marmosets (Liu et al., 2021).

**Suppl. Fig. 8.**
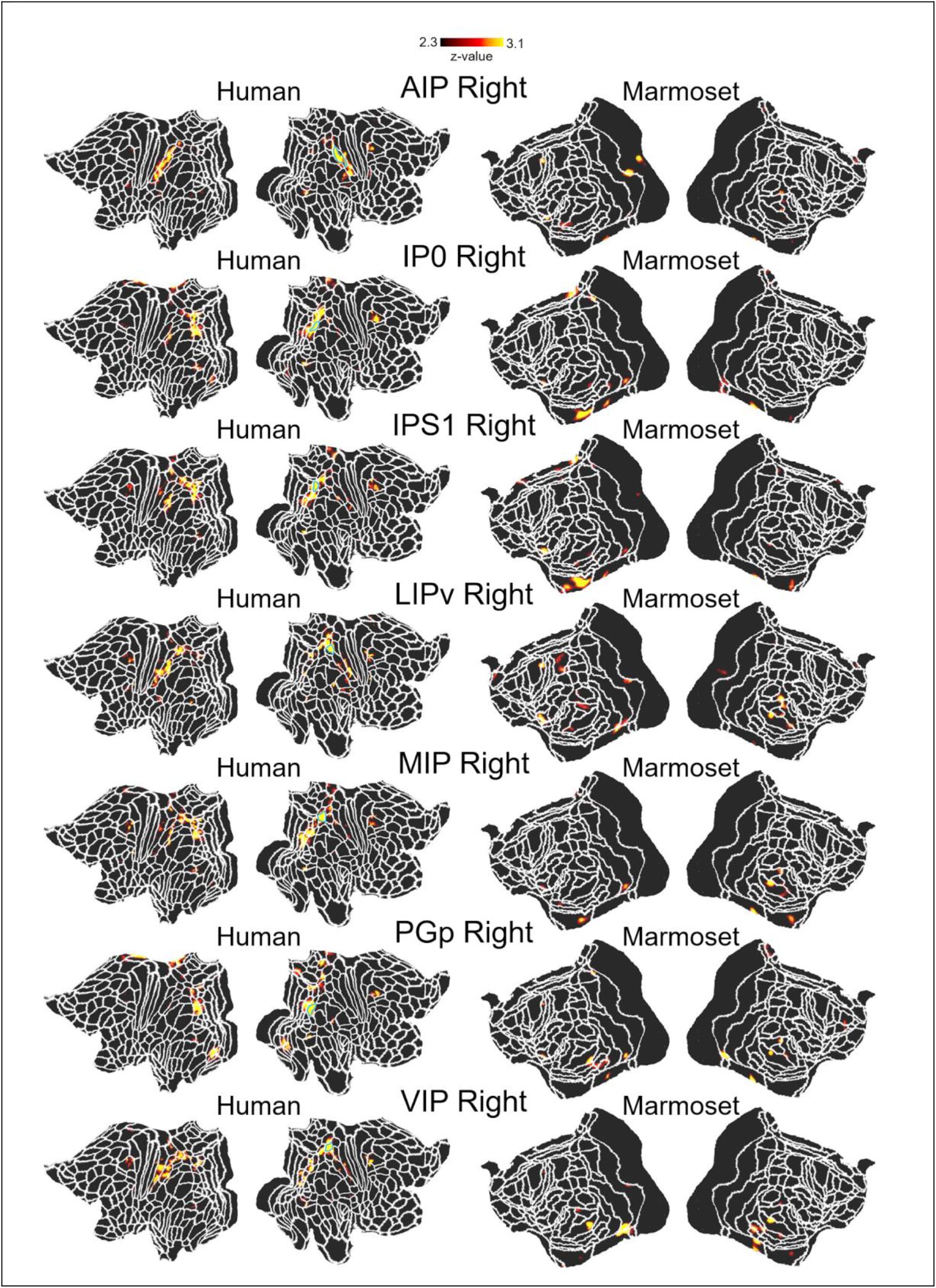
Correlation maps (z-score maps) in human and marmoset brains from each human right parietal volume-of-interest (presented above the map). The maps were shown as thresholded z-score maps on each flattened cortex. The area surrounded by sky blue line indicates the seed regions. The maps for the left VOIs were shown in Suppl. Fig. 7. White lines indicate the borders of the multi-modal cortical parcellation atlas (Glasser et al., 2016) for humans and Paxinos atlas for marmosets (Liu et al., 2021).

**Suppl. Fig. 9.**
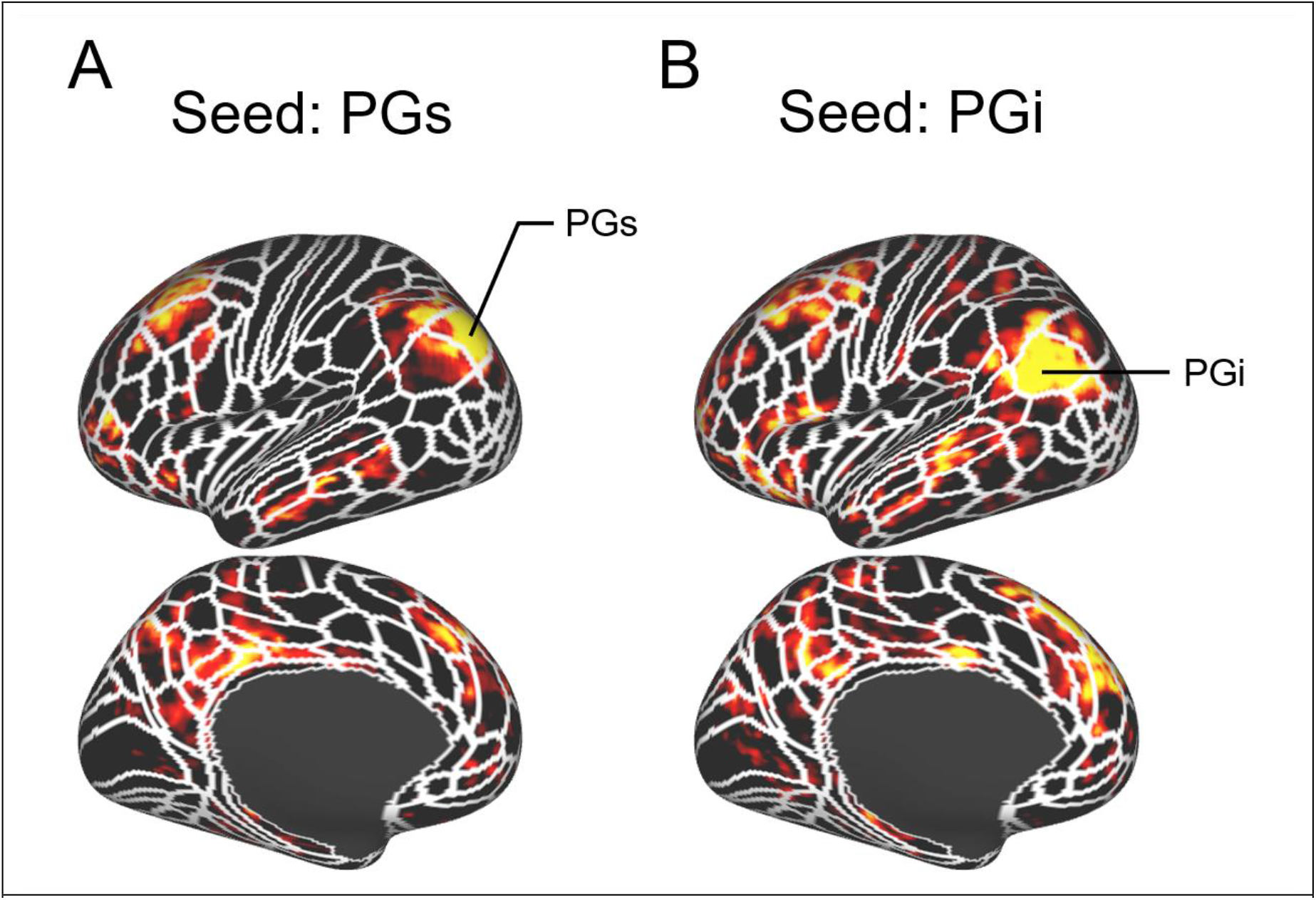
Correlation maps in human brains from each human volume-of-interest (A) PG superior (PGs) and (B) PG inferior (PGi). White lines indicate the borders of the multi-modal cortical parcellation atlas (Glasser et al., 2016) for humans and Paxinos atlas for marmosets (Liu et al., 2021).

